# Successful transfer of myoelectric skill from virtual interface to prosthesis control

**DOI:** 10.1101/2025.08.28.672824

**Authors:** Sigrid Dupan, Simon Stuttaford, Matthew Dyson

**Affiliations:** School of Mechanical and Materials Engineering and School of Electrical and Electronic Engineering, University College Dublin, Ireland; Centre for Biomedical Engineering, University College Dublin, Ireland; Microsystems Group, School of Engineering, Newcastle University, UK

**Keywords:** motor learning, skill transfer, hand prosthesis, electromyography, feedback

## Abstract

**Objective:** Prosthesis control can be seen as a new skill to be learned. To enhance learning, both internal and augmented feedback are exploited. The latter represents external feedback sources that can be designed to enhance learning, e.g. biofeedback. Previous research has shown that augmented feedback protocols can be designed to induce retention by adhering to the guidance hypothesis, but it is not clear yet if that also results in transfer of those skills to prosthesis control. In this study, we test if a training paradigm optimised for retention allows for the transfer of myoelectric skill to prosthesis control.

**Approach:** Twelve limb-intact participants learned a novel myoelectric skill during five one-hour training sessions. To induce retention of the novel myoelectric skill, we used a delayed feedback paradigm. Prosthesis transfer was tested through pre-and post-tests with a prosthesis. Prosthesis control tests included a grasp matching task, the modified box and blocks test, and an object manipulation task, requiring five grasps in total (‘power’, ‘tripod’, ‘pointer’, ‘lateral grip’, and ‘hand open’).

**Main results:** We found that prosthesis control improved significantly following five days of training. Importantly, the prosthesis control metrics were significantly related to the retention metrics during training, but not to the prosthesis performance during the pre-test.

**Significance:** This study shows that transfer of novel, abstract myoelectric control from a computer interface to prosthetic control is possible if the training paradigm is designed to induce retention. These results highlight the importance of approaching myoelectric and prosthetic skills from a skill acquisition standpoint, and open up new avenues for the design of prosthetic training protocols.

## 1 Introduction

Motor learning describes relatively permanent changes in someone’s ability to perform skilled movements [1–3]. During learning, an internal model is built which forms the basis of the feedforward strategy, and feedback is used to reduce the variability in the newly acquired skill [4]. The goal of learning new skills is to perform them in a highly automised and consistent manner. To enhance learning, instructors optimise instructions and feedback, which can be either augmented - also known as extrinsic - or internal [2, 3, 5]. Augmented feedback refers to external or artificially enhanced information, not typically available to the learner during their performance. Whereas, internal feedback refers to the information provided by sensory afferents. To enhance learning, instructors optimise augmented feedback during training. Although internal feedback is present during both learning and eventual skilled performance, augmented feedback tools used during coaching are only provided during the learning phase.

Previous research has shown that frequent augmented feedback can enhance performance during the early acquisition phase of learning, but that these performance gains are lost when feedback is withdrawn [6–8]. This is explained by the guidance hypothesis, which states that permanent feedback during acquisition leads to a dependency on the feedback [9]. One of the reasons Schmidt and colleagues propose for the loss in performance during no-feedback trials, is that guidance overwhelms attention and encourages learners to ignore intrinsic feedback mechanisms during acquisition, even though that may be the only feedback present once training is ‘complete’ [10]. The guidance hypothesis is supported by a range of studies in the fields of motor neuroscience [7, 11–19], sport science [20–22] and physiotherapy [23, 24]. However, reviews show that not all studies reinforce the guidance hypothesis [13, 25, 26], with the differences in results hypothesised to be linked to the complexity of the task [25, 27].

Levels of motor learning can be assessed through short- and long-term retention tests, when augmented feedback is withdrawn, or in transfer tests, when different or related skills are performed [3, 5]. Retention refers to the persistence of the performance, and is therefore considered at the behavioural level. Retention tests are performed a period of time after the training has occurred, while transfer is usually defined as the gain in the ability for performance in one task as the result of practice of another task. These tasks can differ on either the level of the task itself, or through changes in the context or situation the task is performed in [3]. Therefore, retention and transfer tests are very similar. In both cases, the interest is in the persistence of the acquired capability for performance. The two types of tests differ only in that the transfer tests has participants switching to different tasks or conditions, whereas retention tests usually involves retesting participants on the same task or conditions. When transfer tests include an interval of time between the practice and the test itself, skills will need to be internalised and recalled, therefore it follows that retention is a prerequisite requirement to transfer.

Controlling a prosthesis, irrespective of the control method, can be considered a new skill that needs to be acquired. Indeed, a prosthesis can be seen as a new effector for a previously existing behaviour. However, due to the current inability to fully replicate the control of an intact hand, there will inevitably be differences in task demands when using a prosthesis. It follows that myoelectric prosthesis control can be viewed as equivalent to learning to play tennis, either from scratch or when one is already proficient in badminton - depending on the intuitiveness of the control scheme and task demands.

Asymmetric task demands between intact and multi-articulating hands arise in part due to differences in anatomy compared to the human hand, the absence of tactile and proprioceptive feedback, and inaccuracies in decoding movement intentions [28]. This was highlighted in research where a prosthesis was controlled based on kinematic hand data recorded through a 18-DOF data glove. The research showed that even when a prosthesis is controlled based on the kinematics of an intact hand, the prosthetic movement does not fully synchronize with the glove data. Interestingly, when the researchers tracked performance over time, a performance increase of glove-controlled prosthetic movements was observed [28]. This suggests that the participants were using the sensory discrepancy between motor command and task outcome to adapt their input to minimize error. This is a hallmark of motor learning and indicates that even highly biomimetic control will require skill acquisition to a degree.

Historically, prosthetic control has often been treated as an engineering problem rather than a skill acquisition opportunity. For example, early myoelectric devices utilised a threshold-based control [29]. Threshold values were set for the individual upon receiving the device. Producing a contraction that exceeds a threshold requires a relatively coarse motor response compared to a fine control skill that must be learned. Therefore, the nervous system’s capability for producing skilled movement was largely overlooked. Similarly, modern pattern recognition control schemes approach this problem by placing task demands on algorithms rather than the user. The device is adapted to the user’s outputs to make control feel more intuitive. The end-goal of this approach is to create a decoder that perfectly predicts intention, and therefore does not require any adaptation from the user. However, as the research with glove control indicated, perfect synchronization between user and device is challenging - and might never be possible. Therefore, it is likely that a slight misalignment may remain, however small, between user intention and prosthetic device movement. This incongruence ensures that some skill acquisition will be necessary.

The argument that prosthetic control is a skill to be acquired is not solely based on theory. Indeed, when looking at research into long-term use of prosthetic devices, we can see that performance increases [30–35]. As none of these studies included co-adaptation, where the controller changes based on input from the user, the only feasible explanation is learning. Therefore, assuming there are no deliberate changes to the execution of calibration and testing procedures, or to the operation of the device, only two explanations remain for the observed performance gains over time: (1) the user learns to produce *training data* that leads to higher decoding accuracy, or (2) the decoding accuracy of the training data is similar, but the user learns to adapt their *test data*. In both cases, learning is present within the control loop. Therefore, it is necessary to take into account long-standing skill acquisition principles, or risk leaving performance on the table.

Recently, there has been increased interest in prosthetic training, especially through the introduction of myogames, computer interfaces that are controlled by muscle activation. One of the main benefits of myogames is that they can be introduced in the period between amputation and receiving a wearable prosthesis [36, 37]. Due to logistical reasons such as socket manufacturing as well as post-surgery healing, long periods of inactivity risk muscle atrophy. The goal of myogames is to teach people the control necessary for future prosthetic use, with the additional benefit of reducing muscle atrophy. Due to the nature of myoelectric training, where the skills acquired during computer interface interaction must ultimately be applied to the control of a new device, myogames can only be effective if the skills learned during the game transfer to prosthetic control [37].

Research in myoelectric- to prosthesis control transfer has showed varying results [36, 38–44]. Previous work has hypothesized that (1) training with myogames with limited task similarity to prosthesis control led to limited transfer, and (2) providing continuous augmented feedback during training was important for transfer to occur, independent of task similarity. However, subsequent research has shown that transfer can vary for myogames with high task similarity. For example, a mode-switching task led to successful transfer [40], while an machine learning-based task did not [41]. Interestingly, no research has tested for retention prior to testing for transfer. As retention is a prerequisite for transfer, the guidance hypothesis indicates that the common use of realtime feedback in myogames - a type of augmented feedback that is not present in actual prosthetic control - might have a negative effect on retention, and therefore eventual transfer.

We previously demonstrated that the development of a training protocol based on the guidance hypothesis leads to novel myoelectric skills that can be retained [45, 46]. The focus of this study is to test if the presence of retention leads to transfer of these myoelectric skills to prosthetic control. The study follows a pre-post design, where participants completed a range of prosthetic tasks on the first and last day of the study, and completed five 1-hour sessions of training in between. We show that participants significantly improve in several prosthesis control metrics following five days of training. Critically, we show that the training scores participants achieve on trials where augmented feedback was not presented in real time are correlated to prosthetic control ability.

## 2 Methods

### 2.1 Participants

Twelve limb-intact participants (4 female, 8 male) free from neurological or motor disorders volunteered for this study. Ethical approval was granted by the local committee at Newcastle University (Ref: 20-DYS-050). All participants provided written informed consent before the start of the experiment. The overall experiment consisted of two main parts: (1) pre- and post-testing with a prosthesis, and (2) myoelectric training without a prosthesis. Participants completed 5 days of training, with each session lasting 1 hour. Pre-test, training sessions and post-test were all performed on consecutive work days. No participants started the study on a Friday, resulting in all participants receiving one 2-day break during the weekend.

### 2.2 EMG Recordings

Surface EMG signals were recorded with two-channel Gravity analog EMG sensors (OYMotion Technologies Co. Ltd Shanghai, China), which were placed over the flexor carpi radialis (FCR) and extensor carpi radialis (ECR) muscle groups. Signals were acquired using a custom network-enabled myoelectric platform [47], enabling streaming of EMG data over Bluetooth Low Energy to a PC running the AxoPy Python library [48]. Signals were sampled at 500Hz and band-pass filtered between 20 Hz and 150 Hz. Muscle activity was estimated based on the mean absolute value (MAV) over a 760ms sliding window of EMG data, which was updated every 20 ms. Prosthetic control was enabled through the same myoelectric platform. To ensure consistent electrode placement between sessions, the placement of electrodes was indicated with permanent marker on the participant’s skin.

### 2.3 EMG Calibration

A calibration protocol was performed at the start of the pre-test, training period, and post-test, where normalised activity for each channel is:

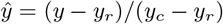

with *ŷ* the normalised muscle activity, y the MAV calculated over the 760ms EMG window, *y_r_* the MAV activity when the participant is at rest, and *y_c_*the MAV activity when the participant performs a comfortable contraction with the muscle targeted by an EMG channel. To determine the comfortable contraction, participants were asked to perform a contraction that they would be able to repeat hundreds of times without fatigue. At the start of the first training sessions, both EMG channels were normalised, and this calibration was used throughout the rest of the training. A separate calibration was performed during the pre-tests as the addition of the weight of the prosthesis was expected to change the calibration. Calibration was repeated during the post-test to take into account the learning participants experienced between the pre- and post-test.

### 2.4 Myoelectric Control Interface

The experimental tasks used during training and prosthetic control are based on the myoelectric control interface (MCI) described in [45, 49], shown in Figure 1a. In the MCI, the position of a 2D circular cursor is controlled by participant’s contractions of the FCR and ECR, where *ŷ* = 1 of each muscle corresponds with the upper limit of it’s corresponding axis. The cursor position reflects the instantaneous *ŷ* of both muscles. Therefore, varying levels of co-contraction allow the cursor to span the entire interface. The control space includes four laterally spaced ‘targets’, and an area corresponding minimal muscle activation referred to as the ‘basket’, Figure 1b. A trial comences once the participant’s muscle activity is considered to be at rest, i.e. when the cursor is positioned within the basket, with 0 *< ŷ <* 0.2. The lower bound of the targets was set at *ŷ* = 0.3, and the top of the target at *ŷ* = 1.

**Figure 1.**
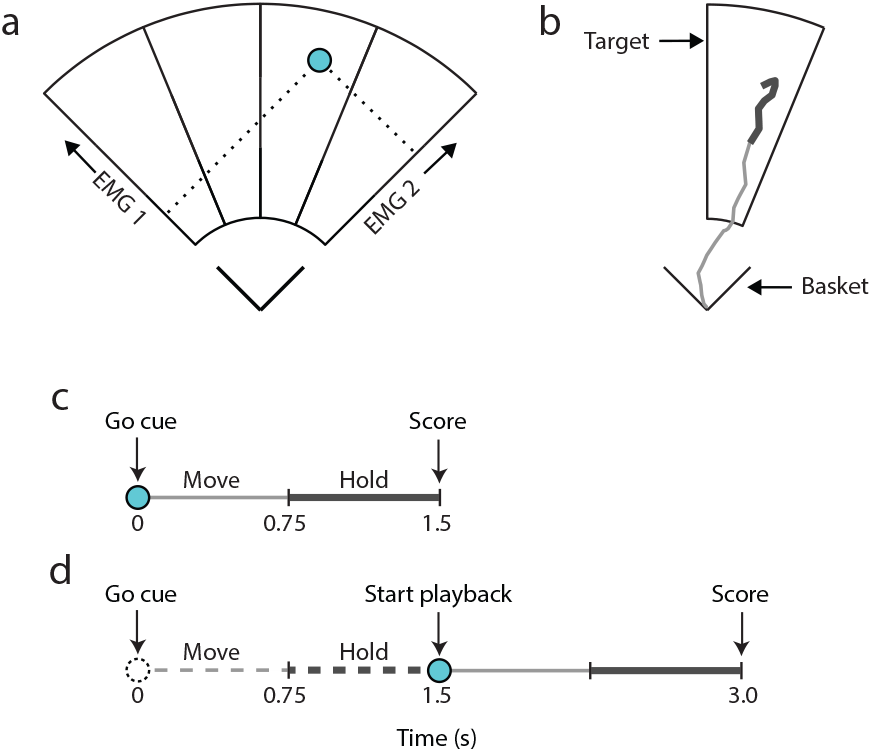
The myoelectric control interface (MCI). (a) The two-dimensional interface space, with the cursor position indicated in blue. (b) An example of a cursor trajectory from basket to target. The thicker line indicates the hold period. (c) Task timing structure for concurrent feedback trials denoting cues, the move and hold periods. (d) Task timing structure for delayed feedback trials denoting cues and the move, hold periods and playback of the cursor’s path. Dashed traces correspond to the ‘blind’ control input window. Solid traces refer to the playback of the cursor’s recorded path during the move and hold periods.

### 2.5 Training Sessions

In between the pre- and post-tests, participants underwent myoelectric training, which was conducted either in their homes or in a laboratory setting. During training, participants completed the MCI task while standing in front of a screen with their elbow flexed at 90*^°^* and their wrist in a neutral position. To initiate a trial, participants had to relax their muscles so that the cursor was within the basket. Once they were at rest, a beep indicated the start of the trial, and one of the four targets was presented. Trials were 1.5s long, and were split into two equal parts: the move and the hold period. The move period ensured participants had enough time to react to and move towards the presented target. At the end of the trial, participants were shown their score, which corresponds with the percentage of time the cursor was within or in contact with the target during the hold period. The scoring mechanism was explained to participants, and they were told to try and maximise their score. Each block contained 60 trials and the targets were presented in a pseudo-random order, ensuring each target was presented 15 times per block.

Experimental blocks contained trials with either concurrent or delayed visual feedback exclusively. When participants received concurrent feedback, the cursor was present during the trial, and reflected the instantaneous normalised muscle activation levels, *ŷ* of both EMG channels. The timing of this condition is shown in Figure 1c. In the delayed feedback condition, the cursor was not visible during the trial. After the completion of the hold period, the cursor’s movement was played back to the participants at the same rate it was captured. This trial structure is illustrated in Figure 1d. The training score was presented at the end of each trial for both feedback conditions. To ensure participants understood the interface and were able to complete all necessary contractions, all participants first experienced a block of concurrent feedback at the start of the first training session. Afterwards, participants were asked to complete as many delayed feedback blocks as they felt comfortable within each session. Participants could request concurrent feedback blocks at any time. Maximising the number of completed delayed feedback blocks was a priority as our previous research showed that delayed feedback facilitated retention, while concurrent feedback blocks did not [45].

### 2.6 Pre- and Post-tests

In both tests, participants wore a transradial bypass socket [50]. The bypass socket is an apparatus that a limb-intact person can wear to simulate prosthetic hand use without the need of a custom prosthesis socket. The bypass socket was fitted with a Touch Bionics robo-limb prosthetic hand and remained attached throughout all tests. As described in section 2.4, the prosthetic hand was controlled through the same myoelectric platform used in training, and the EMG placement was consistent throughout the whole experiment.

#### 2.6.1 Grasp Decoding

Prosthesis control was based on the same MCI described in section 2.4, where each target of the interface was mapped onto one prosthesis grasp. Contrary to the training, trials were not timed during prosthetic control. Instead, trials only commenced once the cursor left the basket, and the grasp was determined by the first target the cursor dwelled within for at least 90% of a 760ms window. Unlike during the training, targets had no upper bound, and thus the user was not penalised for overshooting a target. Once an output grasp was selected, the user could relax and the cursor returned to the basket. No new target would be selected unless the cursor passed through the basket. From left to right, the targets represented the following grips: ‘power’, ‘tripod’, ‘pointer’, and ‘lateral’ grip. The order of the grasps was deliberately selected so that grasps representing finer movements required co-contraction. However, in future clinical applications, any mapping from target to prosthetic grip could be implemented [49,51]. Once the prosthesis was closed, selecting any of the targets resulted in the prosthesis returning to the ‘hand open’ state.

#### 2.6.2 Experimental tasks

The pre- and post-tests consisted of a familiarisation phase and three tasks: grasp matching, box and blocks test, and an object manipulation test.

- **Familiarisation phase**: Participants completed 2 blocks of 60 trials to familiarise themselves with the MCI and calibration of the sensors. In these blocks, targets were presented at the start of each trial, and the participants received concurrent feedback of the cursor based on their muscle activity. Once the cursor was in within or in contact with a target for 750ms, the trial was completed. If the target participants dwelled in was the intended target, a score of 1 was given. Otherwise a score of 0 was recieved. While the participant wore the bypass socket during this task, the prosthesis was not powered on, and therefore did not move.
- **Grasp matching**: Participants were presented with a target on the screen, similar to the familiarisation phase. However, during the grasp matching task, no feedback was presented to the participant, thereby testing the retention of skill [3, 45]. Each participant completed 2 blocks of 60 trails.
- **Box and blocks**: Participants completed a modified version of the box and blocks test [52]. During the test, sixteen wooden cubes were spaced in a 4×4 grid inside a box with a vertical divider (see Figure 2). The goal was to move as many cubes as possible from one side of the box to the other in a given time frame using the prosthesis. Participants were allowed to use any grasp, but were told which grips were most suitable to use (i.e. the power and tripod grasps) and given a practice run of 15 seconds. Each attempt was limited to 60 seconds, and was repeated a total of five times.
- **Object manipulation**: This task required participants to actuate a series of grasps to move a set of objects in a sequence. The objects were evenly spaced on a table in a 2×4 grid marked by tape. The sequence of grasps and movements were: key grip (move pen), point grip (click a mouse button), tripod grip (move screwdriver), power grip (move wooden cylinder). When placing an object, participants would move it to the empty location in the corresponding column marked on the table. Then the participants repeated the tasks in reverse order, which concluded a single trial. Participants were asked to interact with each object with the correct grasp as quickly as possible. If an incorrect grasp was made, the participants had three attempts to repeat the movement. After the third attempt, they were told to move on to the next object.

**Figure 2.**
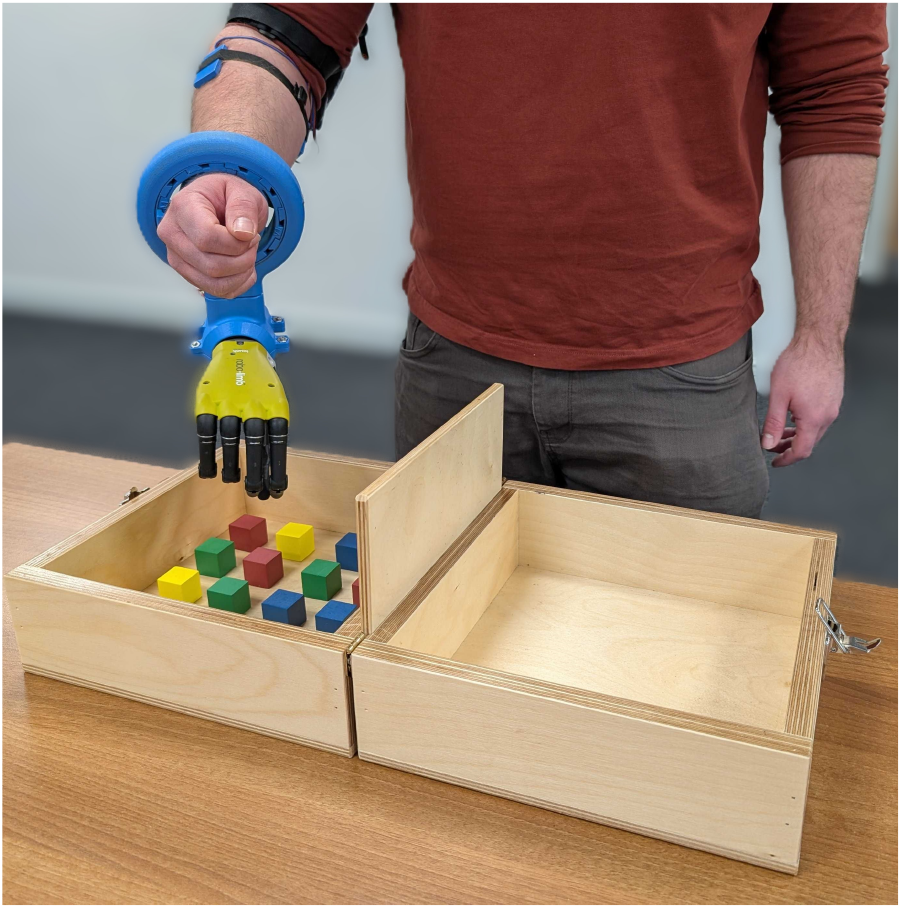
Intact-limb participant use a bypass socket fitted with a prosthetic hand, completing a box and blocks task. Picture is of one of the co-authors demonstrating the task.

### 2.7 Measures and Statistical Analyses

Visual inspection of Q-Q plots suggested that data were not normally distributed. Therefore, non-parametric statistical tests were used. All statistical comparisons were performed using Mann-Whitney U tests, and correlations using Spearman’s rank correlation coefficient. Unless otherwise stated, all values reported represent the median.

#### Training Metrics

The following metrics were used to quantify performance on the MCI task during training.

- **Training score** is defined as the percentage of time of the hold period the target was within or in contact with the target. This score indicates the ability of the participant to stabilise their muscle output at the correct ratio and amplitude, and is important for stable myoelectric control.
- **Path efficiency** is calculated by dividing the cursor’s trajectory by the optimal path length and represents how straight the line of the cursor is. Straight lines, and therefore high path efficiencies, indicate that the ratio of activity between the two EMG sites is internalised and activation of the muscles happens in parallel. Conversely, curved lines are often seen with concurrent feedback when participants adapt the ratio of activities based on the feedback they receive. The end of the cursor’s trajectory was taken to be where the cursor first intersected the correct target. The optimal path length was the distance between the cursor’s starting point and the centroid of the correct target. If the cursor intersected the target below the target centroid, the optimal path end point was taken at the height of the intersection.
- **Coefficient of variation** is a measure of signal variability and is calculated by normalising the standard deviation of a sample of EMG by its mean absolute value. This measure takes into account the nature of EMG signals, which are affected by signal-dependent noise, i.e. the signals become inherently more noisy when they have a higher amplitude. Trials with a low coefficient of variation indicate stable control by the participant. Reported values were calculated from EMG activity during the hold period.

#### Test Metrics

The following metrics were used to quantify performance on the MCI task during prosthesis control.

- **Decoder score:** Performance during the grasp matching task was determined by the decoder score, the percentage of trials in each block where the intended target was successfully decoded.
- **Amount of blocks:** Performance in the box and blocks task was measured by the number of cubes participants were able to transfer in each trial. Only one cube was counted if multiple cubes were deposited with a single grasp.
- **Time:** The time required to complete an object manipulation trial in seconds.
- **Repetitions/trial:** The number of grasp errors during an object manipulation trial.
- **Dropped objects/trial:** The number of times an object was dropped due to unintentional activation of the hand during an object manipulation trial

## 3 Results

### 3.1 Training Analysis

Figure 3 shows an overview of the participants’ mean performance during the delayed feedback training. On average, participants completed a total of 32 delayed feedback blocks, or 1920 trials, over the five days of training. A general trend of improvement can be seen between the first and fifth days (day 1: *M ± std* = 0.60 *±* 0.15; day 5: *M ± std* = 0.75 *±* 0.12).

**Figure 3.**
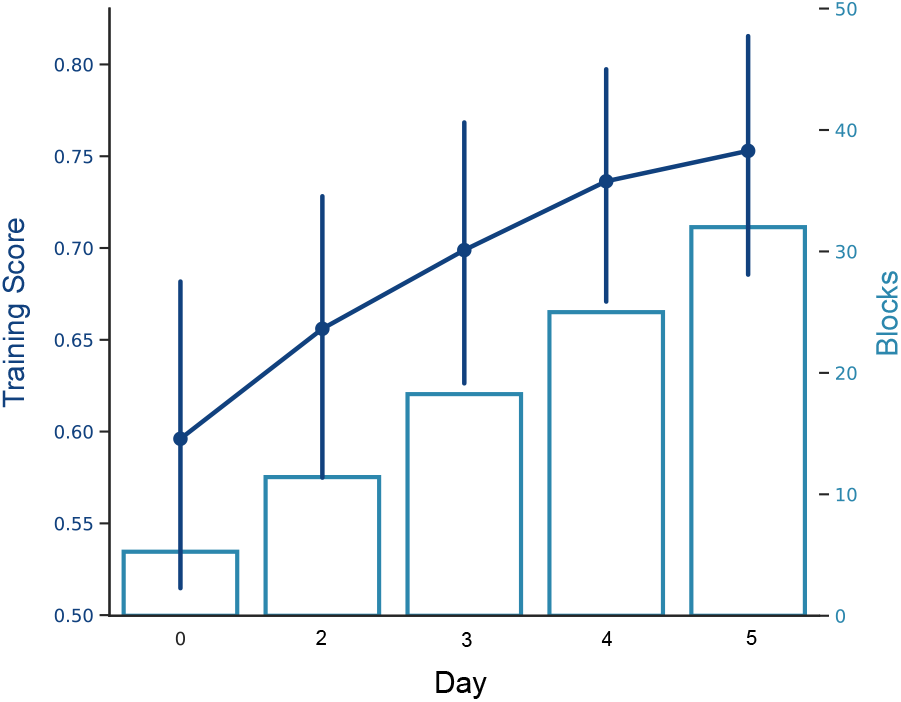
Myoelectric delayed feedback performance during the five day training period. Points, mean scores; error bars, 95% confidence interval; bars, mean number of cumulative blocks completed.

Figure 4 presents evidence that human learning occurs over the training period. Figure 4(a) shows cursor traces for each target from a single participant’s first and best delayed feedback blocks. In general, the cursor traces in the participant’s highest scoring block are less curved and more directly reach the target with greater path efficiency. Figure 4(b) shows that there is a strong negative correlation between training score and path efficiency, the strength of which is also relatively consistent over days. This reiterates that scores increase as path efficiency becomes optimal. Figure 4(c) shows that the coefficient of variation decreased significantly after delayed feedback training (first: *Mdn* = 38%, last: *Mdn* = 28%, *p <* 0.01).

**Figure 4.**
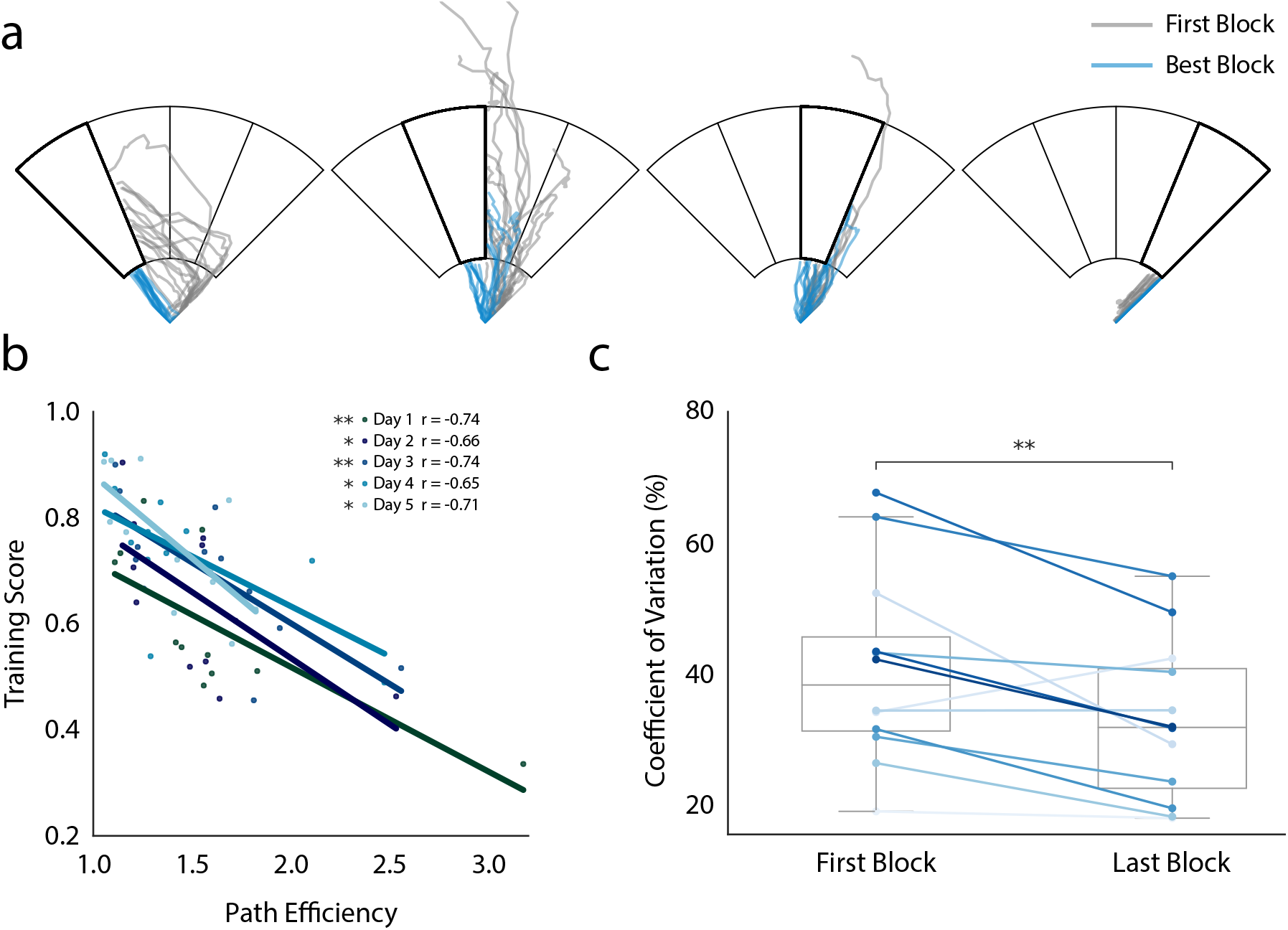
Evidence of human learning driving performance changes. (a) Example of cursor path changes over training. Traces correspond to individual trials from a single participant up until the cursor first contacts the correct target. (b) Relationship between training score and path efficiency over days. Points correspond to participants’ mean score over a single day. (c) Coefficient of variation from each participant’s first and last blocks. Connected points refer to a single participant. Asterisks denote level of significance (∗*, p <* 0.05; ∗∗*, p <* 0.01)

### 3.2 Prosthesis Control Analysis

Figure 5 shows that after training, significant gains in performance occurred for tasks that involved prosthesis control. Improvements were found in the object manipulation task: Figure 5(a) shows the time taken to complete the task significantly decreased (pre: *Mdn* = 116.9 s; post: *Mdn* = 73.4 s; *p <* 0.001), as did the mean number of errors made on a given trial (pre: *Mdn* = 5.0; post: *Mdn* = 2.6; *p <* 0.01), Figure 5(b). In addition, the number of blocks moved in the box and blocks task significantly increased (pre: *Mdn* = 5.4; post: *Mdn* = 7.5; *p <* 0.05), Figure 5(c). The rate of unintentional object drops did not change (*Mdn* = 0).

**Figure 5.**
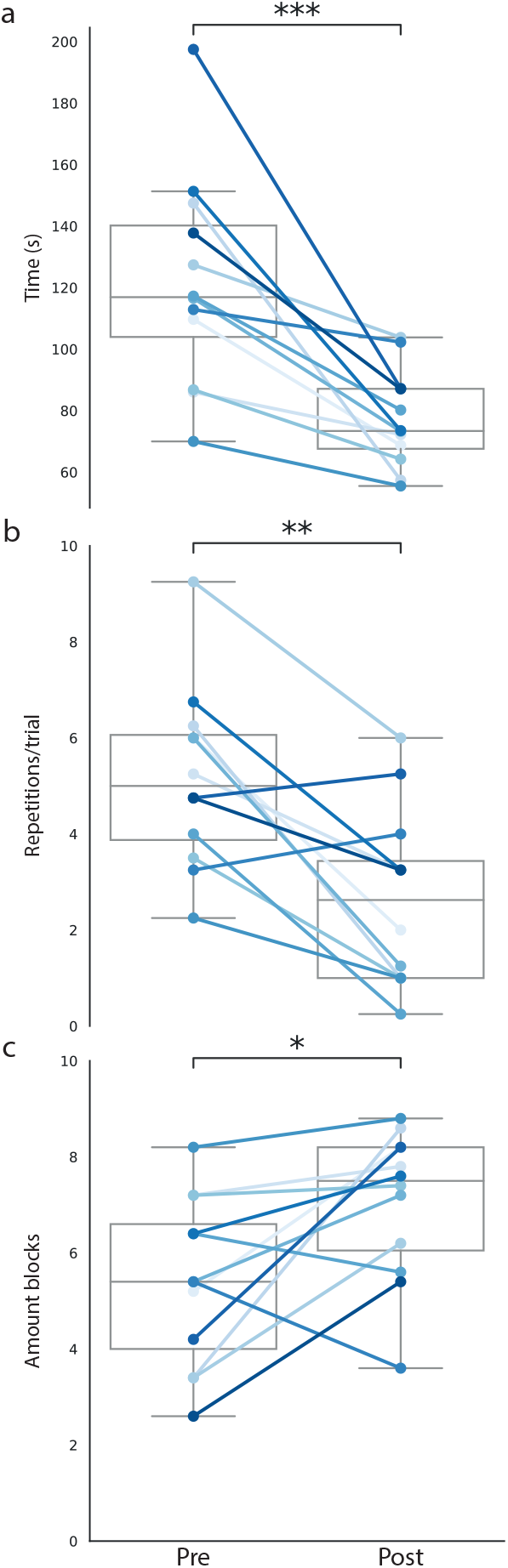
Prosthesis control performance before and after training. (a) Completion time of the object manipulation task. (b) Repetitions during the object manipulation task. (c) Box and blocks task performance. All points correspond to individual participant means. Asterisks denote level of significance. (*, *p <* 0.05; **, *p <* 0.01; *** *p <* 0.001, Wilcoxon signed-rank test)

Analyses were conducted to investigate whether previous experience of the task conditions in the pre-test significantly affected the participants’ performance in the post-test. Figure 6(a-c) illustrates that no relationships between pre- and post-test performance were found for the number of blocks moved during the box and blocks task (*r* = 0.23; *p* = 0.47); the completion time of the object manipulation task (*r* = 0.49; *p* = 0.11); nor the number of errors made during the object manipulation task (*r* = 0.32; *p* = 0.31). In fact, no significant correlations were found between any pre-test metric when compared against itself in the post-test condition.

**Figure 6.**
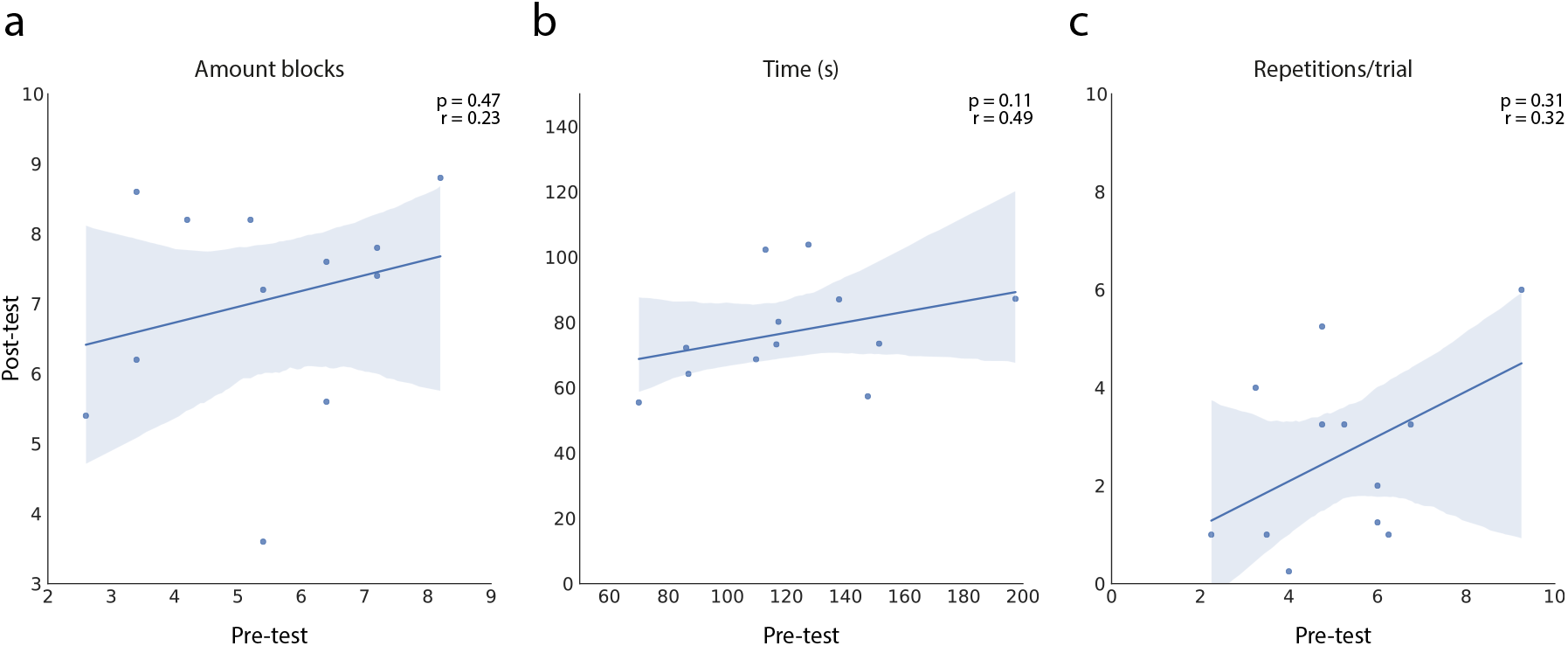
Examination of exposure to previous task conditions. Plots showing no significant correlations between pre-test and post-test performance on the, (a) Box and blocks task; (b) Object manipulation task completion time; (c) Number of repetitions per trial during the object manipulation task.

A positive correlation was found between the mean score achieved during the final day of training and the rate of targets successfully decoded in the post-test (*r* = 0.75; *p <* 0.01), shown in Figure 7(a). Similarly, a moderate negative relationship was found between the final training scores and the mean number of repetitions during the post-test object manipulation task (*r* = −0.63; *p <* 0.05), shown in Figure 7(b). Finally, Figure 7(c) shows no significant relationship between training score and completion time.

**Figure 7.**
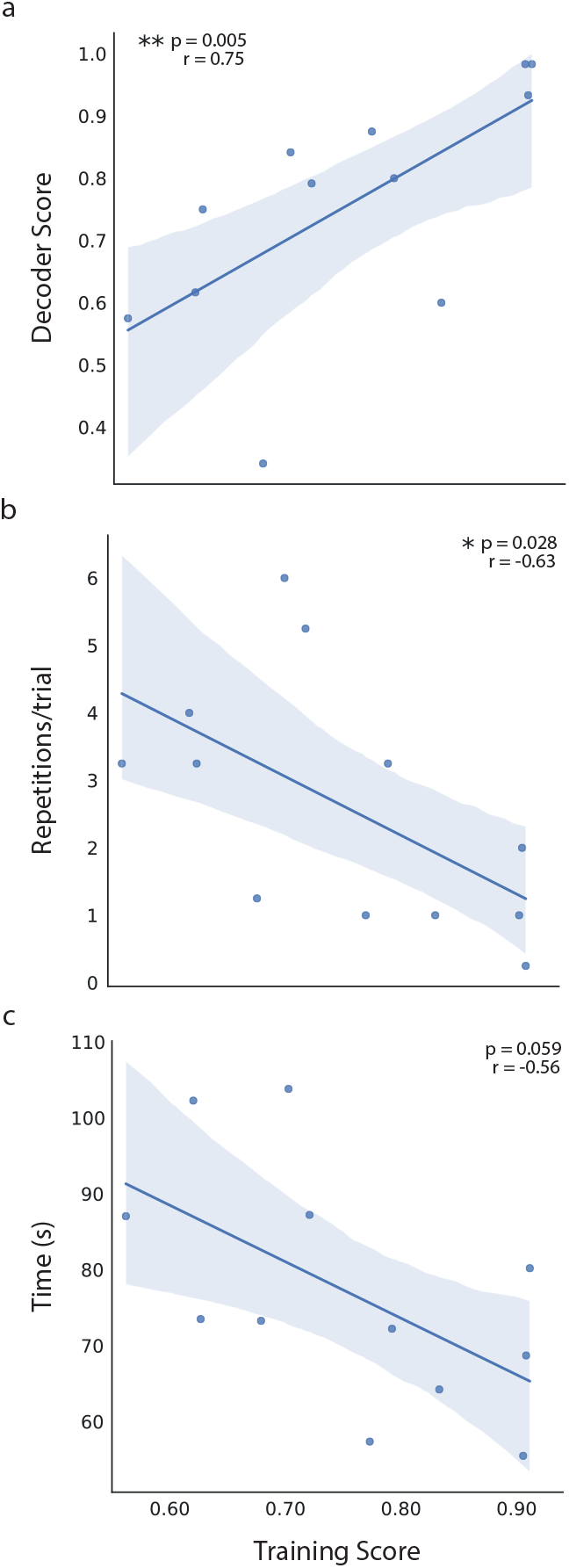
Relationship between myoelectric training and prosthesis control. Correlation between the mean delayed feedback scores from the final day of training to prosthesis control in the post-test. (a) Positive correlation between training performance and decoder score during the zero-feedback condition (*p <* 0.01). (b) Negative correlation between training performance and the mean number of repetitions during the object manipulation task (*p <* 0.05). (c) No significant correlation between training score and the completion time of the object manipulation task (*p* = 0.059). All statistical tests used Spearman’s rank correlation coefficient.

Figure 8 shows the participants’ performance on the grasp matching task in the pre- and post-tests. Although median decoder scores improved between pre- and post-tests (pre: *Mdn* = 0.63; post: *Mdn* = 0.80), the improvement failed to reach significance. In addition, no correlations were found between post-test decoder scores with any other post-test task measures.

**Figure 8.**
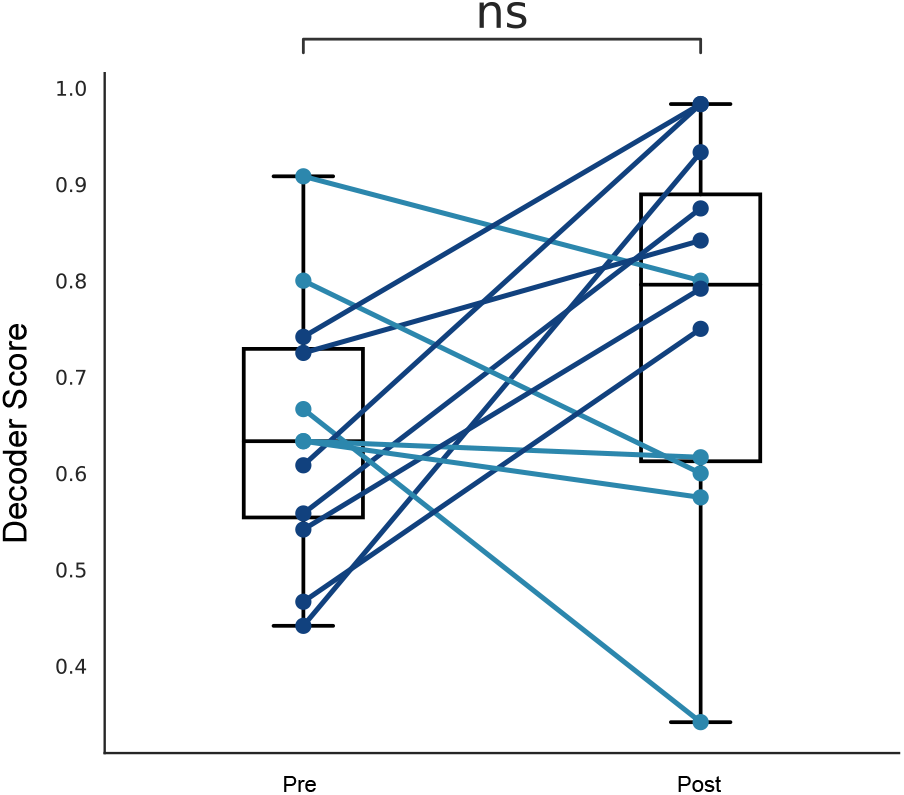
Grasp matching performance during zero-feedback trials in the pre- and post-tests, whilst wearing the prosthesis. Points correspond to individual participant scores. A distinction in line colour is made to visually clarify increases and decreases in decoder scores across individuals.

## 4 Discussion

The current study investigated the retention and transfer of novel myoelectric skills after several days of delayed feedback training. We found that (1) prosthesis control improved following 5 days of training; (2) the prosthesis control metrics were significantly related to the retention metrics during training performance; and (3) increased training results were the result of participants learning and retaining the skill which optimised path efficiency and reduced signal variability. In short, the results of our study suggest that training protocols that induce retention in the training task itself lead to transfer of the skill to prosthetic control.

Since the introduction of the guidance hypothesis in 1984 [6], many strands of research, from fundamental motor control research to applied research in fields such as rehabilitation and sport science, have confirmed that the guiding properties of augmented feedback are beneficial for initial learning, but reduce the ability of the learner to retain the skill [9]. Central to the guidance hypothesis is the idea that augmented feedback feedback not inherent to the learner - can be provided during learning, but is not present during actual performance. Therefore, if initial learning relies on this feedback, retention will not see a similar increase in performance as the feedback the learner is relying on is now withdrawn. Prosthetic training relying on biofeedback, a type of feedback that is currently not present during prosthetic control, should therefore take the guidance hypothesis into account. Our study showed that retention of myoelectric control skills is possible if the training protocol is designed to ensure participants do not rely on concurrent augmented feedback. In our case, this was ensured by introducing delayed feedback [45].

Performance in the delayed feedback training was slightly better than a previous - more structured - delayed feedback protocol [45], even though participants in this study were allowed to introduce concurrent feedback blocks and/or stop the session whenever they wanted. However, the performance was reduced compared to a home-based study in the same publication [45]. These differences can be explained through the general variability in the participant population. Indeed, participants in the current study reached training scores between 0.60 and 0.90 on the final day of training. Additional to the general variability, other factors that could have influenced these scores are engagement, motivation and attention [53, 54]. The same variability in results was also seen in the pre- and post-tests. Interestingly, the performance in the post-tests was correlated to that in the training, while pre- and post-test performance metrics did not show any correlation. As such, the variability included in the participants confirms the hypothesised link between retention and transfer: the more retention you can induce in training, the better the results in the transfer test.

Our study showed an improvement in prosthetic control for both the modified box and blocks task and the object manipulation task, with more significant differences for the object manipulation task. The distinction between the result in both tasks most likely relates to their requirements. While participants can choose the grasp during the box and blocks task, and can therefore complete the task by choosing the grasp that only requires individuation of the flexor (i.e. the power grasp), they have to perform all grasps for the object manipulation task. Previously, we have shown that increases in retention for the MCI task mainly rely on increased performance for the middle targets, which require specific levels of co-contraction [45]. Therefore, increases in transfer performance would mainly be present in tasks that comprise the middle targets rather than those where participants can self-select grasps that are driven by the activation of a single muscle.

The results in this study align with long-standing motor learning literature [7, 8, 12, 14, 15], but are distinct from previous prosthetic research [36, 41–43]. Revisiting the previous research through the lens of the guidance hypothesis might enlighten some of the perceived differences. Previous ‘task-related’ myogames were designed to mimic prosthesis control and/or activities of daily living [40, 55]. By creating a myogame that more closely simulates prosthesis control, similar feedback conditions are inevitably recreated. For example, in van Dijk et al. (2016), a small delay was introduced between the muscle activation and the movement of the virtual gripper [55]. As such, the feedback present during learning resembled that during the actual task. Another paper investigating mode switching did not provide EMG feedback during the virtual switching task [40]. As such, no ‘augmented’ feedback - feedback not present during the actual task - was provided. By contrast, traditional myogames that typically present EMG signals in real time have a greater guiding effect on performance. Therefore, participants may become dependent on the feedback, which leads to low skill retention and transfer. Within this perspective, the similarity of the task goals are less important; instead greater priority is given to the feedback conditions under which the task is performed. Although the level of task similarity is likely key for learning the cognitive mapping between control input and prosthesis response [55], this study shows that transfer without task similarity is possible if feedback similarity is established in training and testing tasks.

Interestingly, despite significant improvements in prosthesis control speed and accuracy, as shown in Figure 5, no corresponding significant improvement was found in post-test grasp matching (Figure 8). Our hypothesis is that the structure of the post-test led to this metric becoming a poor reflection of transfer. Over the five days of training, participants likely became accustomed to completing the task without the additional weight of the bypass socket. Upon commencing the post-test, participants were given two blocks of concurrent feedback trials to check the calibration before completing two blocks of zero-feedback trials. In retrospect, this was a poor experimental choice. Participants may have unknowingly reacted to the visual updates of the cursor and adapted their input on the fly to accommodate a slight shift in calibration, which was likely caused by the weight of the prosthesis.

Ideally, participants should have experienced delayed feedback blocks, instead of concurrent ones, to assess whether the calibration was optimal. This stresses the importance of recreating the feedback loops during transfer tasks, which are similar to those available during real prosthesis control. Although performance over zero-feedback trials was previously shown to be a good estimate of retention [45], the feedback loop in these trials is much more limited compared to real prosthesis control. Therefore, this metric may not have been a good indicator of transfer performance.

The main limitation of the current study was the lack of a control group. This makes it challenging to infer whether performance improvements were due to the training protocol or were simply a result of the participants’ previous experience from completing the pre-test. However, no correlation was found between the pre- and post-test, while the performance on the last training day was significantly correlated to the performance in the post-test. This suggests that the increase in performance is most likely a result of the retention induced during the training instead of the experience during the pre-test. Nevertheless, in the worst-case scenario that it is, in fact, the prior exposure to the pre-test that leads to significant improvements in post-tests which are 7 days apart, that would be an exciting result that has major implications for abstract control! Additionally, this study did not include participants with limb difference. However, this study investigated motor learning and the transfer of skill, and to the best of the authors’ knowledge, there is no indication in any previous research that these central processes would be different in people with limb difference. As this experiment was performed over 7 working days, and no assurance of success, we made the concious choice not to include participants with limb difference.

## 5 Conclusion

In this study, we showed that skills acquired during an abstract training task - specifically designed to induce retention - were transferred from a myoelectric computer interface task to prosthetic control. Additionally, we showed that the level of retention during training was correlated with the amount of transfer. This underlines the importance of seeing myoelectric and prosthetic control skills through the lense of skill acquisition and the importance of the guidance hypothesis. Within this perspective, the similarity of training and task goals are less important; instead, greater importance is given to the feedback conditions under which the task is performed. As current prosthetic control has delayed endpoint feedback, this suggests myogames for training could either mimic current control or be designed specifically to align with the guidance hypothesis, e.g. by delaying feedback as shown in this work.

## Data availability statement

The data that supports the findings of this study are available upon reasonable request from the authors.

## Acknowledgements

This work has emanated from research jointly funded by Taighde Éireann - Research Ireland under Grant number 21/PATH-S/9605, and by the Engineering and Physical Sciences Research Council (EPSRC) via grant EP/R511584/1.

## References

[1] Daniel B Willingham. A neuropsychological theory of motor skill learning. Psychological review, 105(3):558, 1998.

[2] Richard Magill and David I Anderson. Motor learning and control. McGraw-Hill Publishing New York, 2010.

[3] R. A. Schmidt, T. D. Lee, C. Winstein, G. Wulf, and H. N. Zelaznik. Motor Learning Concepts and Research Methods. Human Kinetics, Champaign, IL, 6 edition, 2019.

[4] L. Shmuelof, J. W. Krakauer, and P. Mazzoni. How is a motor skill learned? change and invariance at the levels of task success and trajectory control. Journal of neurophysiology, 108(2):578–594, 2012.

[5] Donna M Bowers, Andrea Oberlander, Kevin K Chui, Kimberly Leigh Malin, and Michelle M Lusardi. Motor control, motor learning, and neural plasticity in orthotic and prosthetic rehabilitation. In Orthotics & Prosthetics in Rehabilitation 4th edition. 2019.

[6] A. W. Salmoni, R. A Schmidt, and C. B. Walter. Knowledge of results and motor learning: a review and critical reappraisal. Psychol. Bull., 95(3):355–386, 1984.

[7] C. J. Winstein and R. A. Schmidt. Reduced frequency of knowledge of results enhances motor skill learning. J. Exp. Psychol. Learn. Mem. Cogn., 16(4):677–691, 1990.

[8] Dana Maslovat, Kirstin M Brunke, Romeo Chua, and Ian M Franks. Feedback effects on learning a novel bimanual coordination pattern: support for the guidance hypothesis. Journal of motor behavior, 41(1):45–54, 2009.

[9] R. A Schmidt. Frequent augmented feedback can degrade learning: Evidence and interpretations. Tutorials in Motor Neuroscience, pages 59–75, 1991.

[10] R. A Schmidt and G. Wulf. Continuous concurrent feedback degrades skill learning: Implications for training and simulation. Hum. Factors, 39(4):509–525, 1997.

[11] Richard A Schmidt, Douglas E Young, Stephan Swinnen, and Diane C Shapiro. Summary knowledge of results for skill acquisition: support for the guidance hypothesis. Journal of Experimental Psychology: Learning, Memory, and Cognition, 15(2):352, 1989.

[12] Pierre-Michel Bernier, Romeo Chua, and Ian M Franks. Is proprioception calibrated during visually guided movements? Experimental brain research, 167:292–296, 2005.

[13] Franz Marschall, Andreas Bund, and Josef Wiemeyer. Does frequent augmented feedback really degrade learning? a meta-analysis. Bewegung und Training, 1, 2007.

[14] Herbert Heuer and Mathias Hegele. Constraints on visuo-motor adaptation depend on the type of visual feedback during practice. Experimental Brain Research, 185(1):101–110, 2008.

[15] Sandra Sülzenbrück and Herbert Heuer. Type of visual feedback during practice influences the precision of the acquired internal model of a complex visuo-motor transformation. Ergonomics, 54(1):34–46, 2011.

[16] Renaud Ronsse, Veerle Puttemans, James P Coxon, Daniel J Goble, Johan Wagemans, Nicole Wenderoth, and Stephan P Swinnen. Motor learning with augmented feedback: modality-dependent behavioral and neural consequences. Cerebral cortex, 21(6):1283–1294, 2011.

[17] Shailesh S Kantak and Carolee J Winstein. Learning–performance distinction and memory processes for motor skills: A focused review and perspective. Behavioural brain research, 228(1):219–231, 2012.

[18] Mohammad Hossein Zamani and Mehdi Zarghami. Effects of frequency of feedback on the learning of motor skill in preschool children. International Journal of School Health, 2(1):1–6, 2015.

[19] Yoichiro Aoyagi, Eri Ohnishi, Yoshinori Yamamoto, Naoki Kado, Toshiaki Suzuki, Hitoshi Ohnishi, Nozomi Hokimoto, and Naomi Fukaya. Feedback protocol of ‘fading knowledge of results’ is effective for prolonging motor learning retention. Journal of Physical Therapy Science, 31(8):687–691, 2019.

[20] Peter JK Smith, Stephen J Taylor, and Keith Withers. Applying bandwidth feedback scheduling to a golf shot. Research Quarterly for Exercise and Sport, 68(3):215–221, 1997.

[21] Douglas L Weeks and Raymond N Kordus. Relative frequency of knowledge of performance and motor skill learning. Research Quarterly for Exercise and Sport, 69(3):224–230, 1998.

[22] Nir Dov Shimony, Ronnie Lidor, and Gal Ziv. The effectiveness of bandwidth knowledge of results on a throwing task in goalball players with visual impairments. European Journal of Adapted Physical Activity, 13(2), 2020.

[23] Carolee J Winstein, Patricia S Pohl, and Rebecca Lewthwaite. Effects of physical guidance and knowledge of results on motor learning: support for the guidance hypothesis. Research quarterly for exercise and sport, 65(4):316–323, 1994.

[24] SMP Verschueren, SP Swinnen, René Dom, and Willy De Weerdt. Interlimb coordination in patients with parkinson’s disease: motor learning deficits and the importance of augmented information feedback. Experimental brain research, 113:497–508, 1997.

[25] Roland Sigrist, Georg Rauter, Robert Riener, and Peter Wolf. Augmented visual, auditory, haptic, and multimodal feedback in motor learning: a review. Psychonomic bulletin & review, 20:21–53, 2013.

[26] Emma Louise Petancevski, Joshua Inns, Job Fransen, and Franco Milko Impellizzeri. The effect of augmented feedback on the performance and learning of gross motor and sport-specific skills: A systematic review. Psychology of sport and exercise, 63:102277, 2022.

[27] Gabriele Wulf and Charles H Shea. Principles derived from the study of simple skills do not generalize to complex skill learning. Psychonomic bulletin & review, 9(2):185–211, 2002.

[28] A. Krasoulis, S. Vijayakumar, and K. Nazarpour. Effect of user practice on prosthetic finger control with an intuitive myoelectric decoder. Frontiers in neuroscience, 13:891, 2019.

[29] A. D. Roche, H. Rehbaum, D. Farina, and O. C. Aszmann. Prosthetic myoelectric control strategies: a clinical perspective. Current Surgery Reports, 2:1–11, 2014.

[30] G. C. Matrone, C. Cipriani, M. Chiara Carrozza, and G. Magenes. Real-time myoelectric control of a multi-fingered hand prosthesis using principal components analysis. J. Neuroeng. Rehabilitation., 9(40), 2012.

[31] H. Bouwsema, C. K. van der Sluis, and R. M. Bongers. Changes in performance over time while learning to use a myoelectric prosthesis. J. Neuroeng. Rehabilitation., 11(16), 2014.

[32] Asim Waris, Irene Mendez, Kevin Englehart, Winnie Jensen, and Ernest Nlandu Kamavuako. On the robustness of real-time myoelectric control investigations: A multiday Fitts’ law approach. J. Neural Eng., 16(2), 2019.

[33] Xinjun Sheng, Bo Lv, Weichao Guo, and Xiangyang Zhu. Common spatial-spectral analysis of EMG signals for multiday and multiuser myoelectric interface. Biomedical Signal Processing and Control, 53, 2019.

[34] M. B. Kristoffersen, A. W. Franzke, C. K. van der Sluis, A. Murgia, and R. M. Bongers. The effect of feedback during training sessions on learning pattern-recognition-based prosthesis control. IEEE Trans. Neural Syst. Rehabilitation Eng., 27(10):2087–2096, 2019.

[35] L. J. Hargrove, L. A. Miller, K. Turner, and T. A. Kuiken. Myoelectric pattern recognition outperforms direct control for transhumeral amputees with targeted muscle reinnervation: A randomized clinical trial. Sci. Rep., 7:13840, 2017.

[36] L. van Dijk, C. K. van der Sluis, H. W. van Dijk, and R. M. Bongers. Learning an EMG controlled game: Task specific adaptations and transfer. PLoS One, 11(8):e0160817, 2016.

[37] C. A. Garske, M. Dyson, S. Dupan, G. Morgan, and K. Nazarpour. Serious games are not serious enough for myoelectric prosthetics. JMIR serious games, 9(4):e28079, 2021.

[38] Ludger Van Dijk, Corry K van der Sluis, Hylke W van Dijk, and Raoul M Bongers. Task-oriented gaming for transfer to prosthesis use. IEEE transactions on neural systems and rehabilitation engineering, 24(12):1384–1394, 2015.

[39] Alexander Boschmann, Dorothee Neuhaus, Sarah Vogt, Christian Kaltschmidt, Marco Platzner, and Strahinja Dosen. Immersive augmented reality system for the training of pattern classification control with a myoelectric prosthesis. Journal of neuroengineering and rehabilitation, 18(1):25, 2021.

[40] A. Heerschop, C. K. van der Sluis, and R. M. Bongers. Transfer of mode switching performance: from training to upper-limb prosthesis use. Journal of NeuroEngineering and Rehabilitation, 18(1):1–16, 2021.

[41] M.B. Kristoffersen, A.W. Franzke, R.M. Bongers, M. Wand, A. Murgia, and C.K van der Sluis. User training for machine learning controlled upper limb prostheses: a serious game approach. J. NeuroEng. Rehabil., 18(1):1–15, 2021.

[42] Christopher L Hunt, Yinghe Sun, Shipeng Wang, Ahmed W Shehata, Jacqueline S Hebert, Marlis Gonzalez-Fernandez, Rahul R Kaliki, and Nitish V Thakor. Limb loading enhances skill transfer between augmented and physical reality tasks during limb loss rehabilitation. Journal of NeuroEngineering and Rehabilitation, 20(1):16, 2023.

[43] Bart Maas, Corry K Van Der Sluis, and Raoul M Bongers. Assessing the effectiveness of serious game training designed to assist in upper limb prothesis rehabilitation. Frontiers in Rehabilitation Sciences, 5:1353077, 2024.

[44] Bart Maas, Jack Tchimino, Bram van Dijk, Alessio Murgia, Corry K van der Sluis, and Raoul M Bongers. Skill transfer of upper limb prosthesis control after training in a virtual reality environment. Technology and Disability, 37(2):137–148, 2025.

[45] S. A. Stuttaford, S. S. G. Dupan, K. Nazarpour, and M. Dyson. Delaying feedback during pre-device training facilitates the retention of novel myoelectric skills: a laboratory and home-based study. Journal of Neural Engineering, 20(3):036008, 2023.

[46] S. A. Stuttaford, M. Dyson, K. Nazarpour, and S. S. G. Dupan. Reducing motor variability enhances myoelectric control robustness across untrained limb positions. IEEE Transactions on Neural Systems and Rehabilitation Engineering, 2023.

[47] M. Dyson, J. Olsen, and S. Dupan. A network-enabled myoelectric platform for prototyping research outside of the lab. In 2021 43rd Annual International Conference of the IEEE Engineering in Medicine & Biology Society (EMBC), pages 7422–7425. IEEE, 2021.

[48] K. R. Lyons and B. W. L. Margolis. Axopy: A python library for implementing human-computer interface experiments. Journal of Open Source Software, 4(34):1191, 2019.

[49] M. Dyson, J. Barnes, and K. Nazarpour. Myoelectric control with abstract decoders. J. Neural Eng., 15(5):056003, 2018.

[50] M.D. Paskett, N.R. Olsen, J.A. George, D. T. Kluger, M. R. Brinton, T. S. Davis, C. C. Duncan, and G.A. Clark. A modular transradial bypass socket for surface myoelectric prosthetic control in non-amputees. IEEE Transactions on Neural Systems and Rehabilitation Engineering, 27(10):2070–2076, 2019.

[51] T. Pistohl, C. Cipriani, A. Jackson, and K. Nazarpour. Abstract and proportional myoelectric control for multi-fingered hand prostheses. Ann. Biomed. Eng., 41(12):2687–2698, 2013.

[52] Jacqueline S Hebert, MBA Justin Lewicke, et al. Normative data for modified box and blocks test measuring upper-limb function via motion capture. Journal of rehabilitation research and development, 51(6):919, 2014.

[53] Jan B Engelmann, Eswar Damaraju, Srikanth Padmala, and Luiz Pessoa. Combined effects of attention and motivation on visual task performance: transient and sustained motivational effects. Frontiers in human neuroscience, 3:342, 2009.

[54] Joo-Hyun Song. The role of attention in motor control and learning. Current opinion in psychology, 29:261–265, 2019.

[55] L. Van Dijk, C. K. van der Sluis, H. W. van Dijk, and R. M. Bongers. Task-oriented gaming for transfer to prosthesis use. IEEE Transactions on Neural Systems and Rehabilitation Engineering, 24(12):1384–1394, 2015.

